# *De Novo* Design of Peptide Masks Enables Rapid Generation of Conditionally-Active Miniprotein Binders

**DOI:** 10.1101/2025.02.25.639922

**Authors:** Cristina Montaner, Roberta Lucchi, Montserrat Escobar-Rosales, Marc Expòsit, Cristina Díaz-Perlas, David Baker, Benjamí Oller-Salvia

## Abstract

The widespread expression of many therapeutic targets in both diseased and healthy tissues presents a significant challenge for protein therapeutics, often resulting in dose-limiting side effects. To focus the action of biotherapeutics at the disease site, they can be reversibly inactivated by tethering an affinity mask through a linker cleavable by a disease-specific cue. However, current methods to generate affinity masks require extensive screening of masking sequence libraries. Here, we report a workflow to design the first *de novo* peptide masks for the reversible inactivation of miniprotein binders and we show its application on an EGFR antagonist miniprotein. By extending the C-terminus of the miniprotein to cover the binding interface, a masking sequence is generated that decreases binding to EGFR over 1000 times. Binding of the parental miniprotein can be fully rescued by cleavage of the linker between the binder and the mask with tumor-specific proteases. Additionally, by site-specifically conjugating the peptide mask through a chemical linker, we show we can render the protein responsive to alternative stimuli such as light. Our approach opens the doors to dramatically accelerating the development of affinity-based masks for other proteins with therapeutic potential to render them conditionally active to any stimulus.

## INTRODUCTION

Protein therapeutics are rapidly growing and have contributed to tackling a variety of diseases, ranging from cancer to immune disorders.^[1]^ One of the keys to their success is the high selectivity for their targets. However, most targets are not only present at the diseased site but also in healthy tissues. This may result in dose-limiting side effects. A way to address this issue is by generating protein therapeutics that are only activated at the target site. Many strategies have been developed to render proteins conditionally active to endogenous or exogenous cues.^[2,3]^ One of the most efficient and well-established methods for the reversible inactivation of biologics, such as antibodies and cytokines, is the use of affinity-based masks. These masks interact with the binding site, thereby preventing target engagement while they are not removed. Development of affinity-based masks generally requires labor- and time-intensive screening of protein libraries. Here, we propose the first method that enables the rapid computational design of peptide masks *de novo*.

An affinity-based mask^[4,5]^ typically consists of an N-terminal extension containing an amino acid sequence that interacts with the protein binding site to prevent target engagement. The mask is linked to the protein therapeutic by a cleavable linker sensitive to internal or external cues such as proteases overexpressed in the tumor or light.^[5][6][7][8]^Although affinity-based masks were originally reported for antibodies,^[4]^ their usage has been extended to different types of proteins with therapeutic potential, including immunoglobulins (IgGs), antigen binding fragments (Fabs), affibodies, and cytokines.^[2,9]^ Masking moieties range from linear peptides^[5]^ to chain antibody fragments (scFv),^[6]^ nanobodies,^[7]^ and affibodies.^[8]^ The affinity of the mask for the paratope must be sufficiently high when fused or tethered near the active site to prevent the antibody or therapeutic protein from binding its target, yet low enough to allow it to diffuse away once cleaved and to be displaced by the target.^[10]^ Cropped domains from the antigen including the epitope have been used in very few occasions due to the low stability of the resulting protein and the difficulty to attain the optimal affinity.^[4,11]^ Such affinity fine-tuning involves screening of extensive peptide or protein libraries. Until now, several approaches have been developed based on bacterial and yeast display libraries, all of which require substantial time and effort.^[5,8,12]^

Traditionally, protein therapeutics have been based on natural biomolecules or closely-related derivatives. However, machine learning strategies have recently revolutionized the field of protein computational modelling and design. It is currently possible to model proteins with a precision close to experimental and even to design proteins with therapeutic potential from scratch.^[13–15]^ At least one *de novo* designed protein-based vaccine has been clinically approved and other protein therapeutics have shown great potential in preclinical and clinical settings.^[1,16]^ The design of miniproteins is particularly accessible through computational methods.^[17]^ Miniprotein binders have several advantages over traditional binders derived from natural biomolecules such as immunoglobulins, including small size (down to 7kDa), improved tissue diffusion, enhanced stability, high production yields, and ease of modification and characterization. An increasing number of miniprotein binders have already been reported to engage a plethora of targets,^[17–19]^ many of them with therapeutic potential. For instance, EGFR-targeting miniproteins have been shown to inhibit the signaling of this receptor overexpressed in cancer cells. However, as with most other receptors, EGFR is not only expressed at high levels on tumor cells but also on some healthy tissues. Off-site engagement of EGFR in the healthy epithelium can trigger side effects such as acute skin rash in some patients.^[20]^ Therefore, miniproteins targeting EGFR and other therapeutic receptors would benefit from an additional layer of specificity through reversible inactivation with a mask that is selectively released at the tumor site.^[4,5]^

To bypass extensive library screenings and accelerate the development of affinity-based masks, here we establish a method to design minimal masks *de novo* and show its applicability on a miniprotein binder against EGFR. We utilize a computational workflow based on the machine-learning software RoseTTAFold inpainting, ProteinMPNN, and AlphaFold2 to extend the C-terminus with a protease-cleavable linker and a sequence to cover the binding interface. ^[21]^ Applying several *in silico* filters, we selected and expressed six designs. All masked designs successfully inhibited interaction with the antigen by over two orders of magnitude. Moreover, the masks could readily be cleaved by a tumor-specific metalloprotease fully rescuing the affinity of the unmodified protein. We showed that the best design could successfully inhibit downstream signaling of EGFR only upon protease activation and studied the interaction between the miniprotein and this peptide mask. Finally, through site-specific conjugation of the mask to the miniprotein binder, we demonstrated that we could implement sensitivity to light, thereby expanding the applicability of the masking approach to other stimuli.

## RESULTS AND DISCUSSION

### Design of masked miniprotein binders

To show we could design a masking peptide from scratch, we selected as a model protein a *de novo* designed miniprotein binder (mb) against EGFR domain I previously reported by the Baker laboratory, EGFRn_mb (Figure 1a).^[17]^ EGFRn_mb is highly thermostable and inhibits the downstream signaling of the EGFR receptor. The miniprotein is a three-helix bundle, with two alfa helices mediating the interaction with EGFR, while the third one acts as a linker between the former two and as a scaffold to stabilize them. We hypothesized that we could design a mask that would cover this binding site and sterically hinder binding with EGFR domain I (Figure 1b). As we aimed to design a minimal masking moiety, we envisioned complementing the three helical bundle with a fourth helix linked to one of the termini via a protease-cleavable sequence.

**Figure 1.**
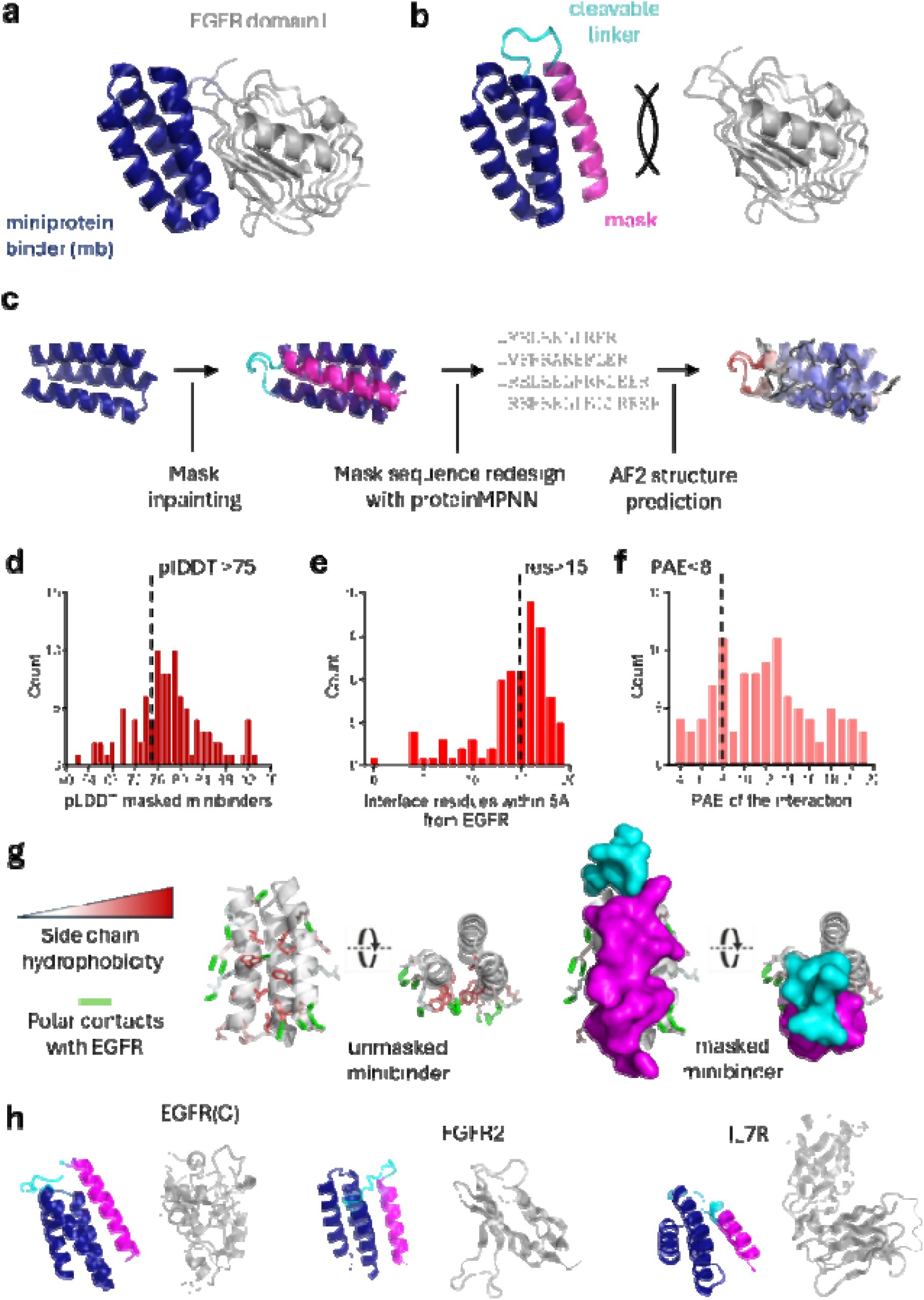
Masked miniprotein binder design. a) Complex of EGFRn_mb miniprotein (deep blue) and its target EGFR domain I (light grey) modelled with AF2.^[17]^ b) Design of an affinity mask (magenta) based on a C-terminal extension of the miniprotein that prevents binding to EGFR and bears a cleavable linker (cyan). c) Workflow of the computational design of the mask. Masked mb structure modelled with AF2 is colored by pLDDT and residues proposed by ProteinMPNN are depicted in black. d) Histogram showing the distribution of pLDDT of masked miniprotein binders. e) Histogram representing the number of residues at 5 Å from EGFR. f) Histogram of the Predicted Alignment Error (PAE). g) EGFRn_mb binding interface with EGFR and mask interference. The 20 residues within 5 Å of the EGFR surface are shown as sticks. Hydrophobicity of the side-chains is indicated in red. Polar contacts between EGFRn_mb and EGFR are depicted as dotted lines in green. The mask is shown in magenta and the linker in cyan. Masked miniproteins in this figure are represented using an AlphaFold3 model of EGFRn_mb with the mask design M3. h) Application of the workflow to generate masks for other miniprotein binders^[17]^ targeting the C-terminal part of EGFR domain I, FGFR2, and IL7Rα shows generalizability of the design strategy.

Given that we aimed to fill a groove between two alpha helices while keeping them in place, we addressed the mask design as a “missing information” recovery problem (Figure 1c). Thus, we elongated one of the termini of the miniprotein binder utilizing RoseTTAFold (RF_joint_) inpainting, a machine-learning model that creates viable protein scaffolds starting from a functional site.^[21]^ We instructed this model to extend the N-or the C-terminus of the miniprotein binder with a fixed protease substrate sequence followed by 15-25 residues, which we anticipated would fold back to interact with the binder. We observed that the C-terminal extensions aligned more naturally with the target groove than N-terminal extensions and decided to focus on the C-terminus. As a linker for the mask, we selected a motif that is recognized by matrix metalloproteinases, mainly MMP2 and MMP9 since they are highly overexpressed in many tumors with high levels of EGFR.^[22]^ All masks would be preceded by a GS linker, and between the miniprotein binder and the protease-cleavable linker, we added 1-3 residues to provide more flexibility to the linker since we reasoned that this could help the protease to recognize efficiently the substrate sequence. With these parameters, 100 designs were generated with RF_joint_ inpainting.

The sequence of the 15-25 “inpainted” residues of the masks was then redesigned using ProteinMPNN.^[23]^ This deep learning model is used to suggest amino acid sequences that encode a target structure. Here, ProteinMPNN provides a sequence for the mask to bind the EGFR binding interface on EGFRn_mb. Other protein design methods such as RFdiffusion can also be utilized in this step by defining the residues to be masked as hotspots (Figure S1). Finally, the structure of the sequences generated was predicted with AlphaFold2 (AF2) and the models obtained were analyzed and filtered to select the best designs for experimental validation. Remarkably, even without providing instructions to the model about the surface to mask, in the vast majority of the designs the mask was naturally led to cover the interaction surface with EGFR. This could be explained given the alpha helical conformation of the surface and the relatively high hydrophobicity of the groove between the two helices involved in binding EGFR. By incorporating the mask in the groove between the two helices present in the 100 designs, all 8 hydrophobic residues between the two helices within 5 Å of EGFR and at least 2 out of 8 residues that appear to establish polar interactions are totally hindered (Figure 1g). Although in this study we focus on EGFRn_mb, similar masks can be obtained for other reported miniprotein binders such as EGFRc, FGFR2, or IL7Rα indicating the wide applicability of our strategy (Figure 1h).

We evaluated the designs based on several *in silico* criteria. For all designs with a reliable structure (pLDDT > 75, Figure 1d and S2),^[15]^ we evaluated the interaction between the mask and the miniprotein binder. To do so, we examined the number of sterically hindered amino acid residues on the miniprotein binder within 5 Å of the mask (Figure 1e), which ranged from 4 to 20. We set a threshold of at least 15 covered residues since a correlation between the number of residues covered and masking capacity should be expected. We added one design (M2, Figure 2a) in which the mask covered only 12 residues to evaluate the relevance of this parameter. To focus on the models with highest accuracy, we established a threshold of 8 for the inter-chain Predicted Aligned Error (PAE) narrowing down the designs to 15 (Figure 1f). Finally, we inspected these designs visually and selected six of them that were diverse in length (Figure 2a and 2b). The models obtained with RF_joint_ inpainting displayed high similarity with the AF2 prediction (RMSD = 1.1±0.2 Å). As we expected from this strategy, all designs display an alpha helical structure forming a four-helix bundle. The helix turns are reflected in all sequences by the amino acid residue pattern that alternates 2-3 polar residues with 1-2 apolar residues (Figure 2a). We evaluated the designs based on several *in silico* criteria. For all designs with a reliable structure (pLDDT > 75, Figure 1d and S2),^[15]^ we evaluated the interaction between the mask and the miniprotein binder. To do so, we examined the number of sterically hindered amino acid residues on the miniprotein binder within 5 Å of the mask (Figure 1e), which ranged from 4 to 20. We set a threshold of at least 15 covered residues since a correlation between the number of residues covered and masking capacity should be expected. We added one design (M2, Figure 2a) in which the mask covered only 12 residues to evaluate the relevance of this parameter. To focus on the models with highest accuracy, we established a threshold of 8 for the inter-chain “Predicted Aligned Error” narrowing down the designs to 15 (Figure 1f). Finally, we inspected these designs visually and selected six of them that were diverse in length (Figure 2a and 2b). The models obtained with RF_joint_ inpainting displayed high similarity with the AF2 prediction (RMSD = 1.1±0.2 Å). As we expected from this strategy, all designs display an alpha helical structure forming a four-helix bundle. The helix turns are reflected in all sequences by the amino acid residue pattern that alternates 2-3 polar residues with 1-2 apolar residues (Figure 2a).

**Figure 2.**
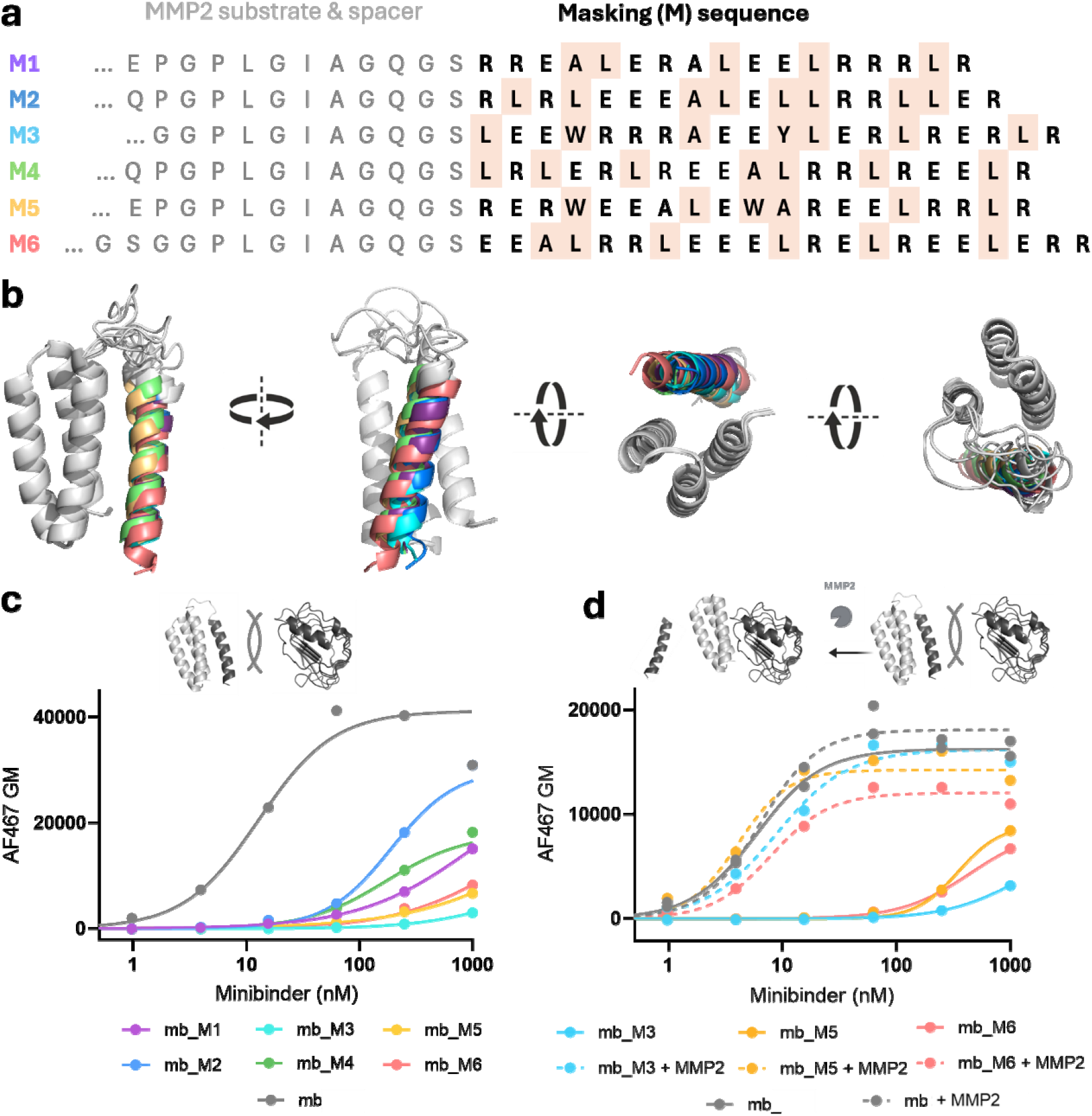
Binding assessment of selected masked miniprotein binders. a) Sequences of the C-terminal extensions of EGFRn_mb (mb) in mb_M1 to mb_M6. Masking sequences binding the miniprotein are on the right in bold. Non-polar residues highlighted in wheat color indicate an alpha helical pattern. The sequence recognized and cleaved by MMP2 and the flanking spacer residues are shown in grey. b) Superimposed predicted structures of the masked mb_M1 to mb_M6 constructs. c) Flow cytometry analysis of the binding assay with the masked miniprotein binders shows reduced affinity compared to the parental unmodified miniprotein. d) Flow cytometry analysis of the binding assay before and after activation with the protease MMP2 shows full recovery of binding affinity.

### Blocking capacity of the masks and affinity recovery upon protease cleavage

The six selected designs and the unmasked miniprotein binder control were cloned and expressed in *E. coli* BL21(DE3) cells in an IPTG-induced pET29 vector. All the designs were produced with an N-terminal 10xHis-tag, that was used for both affinity purification and downstream detection. All miniproteins were obtained with the expected molecular weight and high purity (>90%) as shown by SDS-PAGE (Figure S3) and LC-UV-MS (Figure S4). The yield of purified protein per liter of culture was 78 mg for the unmodified binder and between 18 and 91 mg for the masked designs, indicating that the designed masks did not have a dramatic impact on the miniprotein expression.

The binding capacity of each design was assessed on A-431 epidermoid carcinoma cells that naturally express very high levels of EGFR.^[24]^ Incubation with the cells was performed at 4ºC and binding was assessed via flow cytometry. In all experiments, the EC50 for EGFRn_mb was close to the one reported (6-9 nM vs 1.2 nM). All six designs showed dramatically reduced binding affinity compared to the unmasked EGFRn_mb (mb) (Figure 2c). The increase in EC50 due to the presence of the mask ranged from 20 for the worst design to over 100-fold for the other 5 designs. As expected, the design with lowest inactivation capacity, M2, corresponded to the one with the least number of amino acid residues involved in EGFR binding: 12 aa in M2 vs 16 aa or more in the rest. Although measuring the EC_50_ for these six masked designs was impossible over the concentration ranges tested (from 2 µM to 0.2 nM), at least for the M3 mask and potentially for M5 and M6 the decrease in binding affinity was of over 3 orders of magnitude. This is comparable to state-of-the art affinity masks identified using extensive yeast and bacterial display library screening campaigns^[6,7,12,25]^ and 1-2 orders of magnitude higher than most masking methods based on steric hindrance.^[26–28]^

After having confirmed that the designed masks were able to inhibit the binding of the EGFRn_mb to cells overexpressing the EGFR, we proceeded to evaluate whether protease-mediated cleavage of the linker between the mask and the miniprotein binder could recover binding affinity. To this end, we selected the three designs with the most efficient masks, M3, M5 and M6. We incubated the masked miniproteins with pre-activated MMP2 for 2h at 37 °C, as previously reported.^[29]^ Hydrolysis of the linker was verified by SDS-PAGE and LC-UV-MS (Figure S5 and S6). All three designs were efficiently cleaved, indicating that the protease substrate was accessible in the three designs regardless of the flanking amino acid residues (Figure 2a). Masked miniprotein variants with or without pre-cleavage with MMP2 were incubated with A-431 cells at 4°C. In all three cases, MMP2-mediated cleavage was enough to enable full binding recovery, with the activated miniproteins showing affinities similar to the unmasked control (Figure 2d). This indicates that the mask has sufficient affinity to efficiently block binding when covalently attached to the miniprotein and that, upon cleavage, it may diffuse away and is completely outcompeted by the target antigen, EGFR. These results prove for the first time that *de novo* designed peptide affinity masks can provide effective blockage and conditional activation. The fact that all expressed designs display this behavior is even more remarkable taking into account that the masks were designed on the predicted structure of the anti-EGFR miniprotein binder.

### Functional verification of the lead candidate mb_M3

The M3 mask design consistently provided the best blocking capacity. Therefore, we focused on this design for further experiments. We confirmed that binding upon activation was specific for cells with high levels of EGFR and negligible in cells with low levels of the receptor (Figure 3a and 3b). In addition, absence of binding of mb_M3 and recovery of nanomolar binding affinity upon mask cleavage was confirmed with the isolated receptor using biolayer interferometry (Figure 3c).

**Figure 3.**
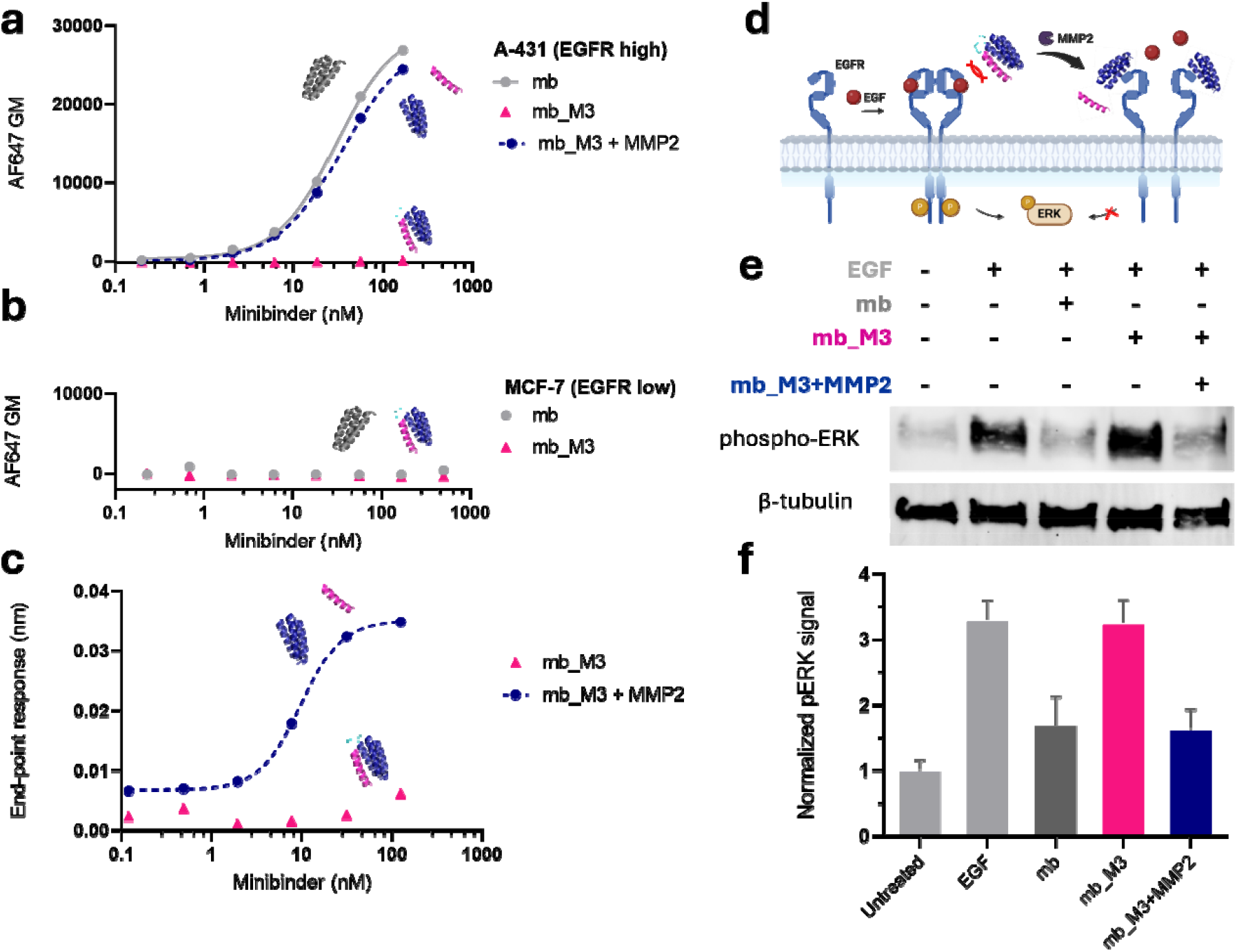
Functional characterization of the lead candidate mb_M3. Flow cytometry analysis of mb_M3 binding, before and after activation, to cells expressing different levels of EGFR: (a) A-431 cells, which express high levels of EGFR, and (b) MCF-7, which express low levels of EGFR. c) Binding of mb_M3 to EGFR before and after proteolytic activation with MMP2 as measured by biolayer interferometry. d) Schematic representation of the EGFR antagonist assay with masked miniprotein binders. e) Western Blot analysis of EGFR antagonism assay shows: basal levels of ERK phosphorylation (lane 1), high phospho-ERK levels upon stimulation with EGF (lane 2), inhibition of EGFR signaling with EGFR_n (mb) (lane 3), no EGFR signaling inhibition with mb_M3 (lane 4), and full rescue of inhibitory capacity upon MMP2 cleavage (lane 5). f) Densitometric quantitative analysis of Western Blot (error bars represent the standard deviation, n = 3).

After the binding assessment, we challenged the capacity of the miniprotein binder to conditionally inhibit EGFR signaling.^[30]^ To this end, we set up an assay in which we activated the MAPK cascade through stimulation with EGF in the presence or absence of the miniprotein binder, which has been shown to be an antagonist of EGFR (Figure 3d).^[17]^ We selected U-87, a glioma-derived cell line, as an example of tumor cells that express high levels of non-mutated EGFR and that have previously been used for this assay.^[31,32]^ In this experiment, the level of downstream phosphorylated ERK (phospho-ERK) was increased (Figure 3e) by incubating these cells with EGF.

We then confirmed that binding of EGF could be efficiently inhibited by the unmodified EGFRn_mb (mb) at two different concentrations (Figure 3e, S7 and S8). As expected, when mb_M3 was incubated, no decrease in phospho-ERK was observed. Conversely, when the masked miniprotein binder was preincubated with the MMP2 protease, it was able to outcompete EGF at the same level as the mb (Figure 3f). These results further prove that the function of the miniprotein binder can be reversibly inactivated with our mask design and reactivated in the presence of tumor-specific proteases.

### Study of mask affinity

In the AlphaFold2 and AlphaFold3 predicted structures of the miniprotein binder and mask M3, there is one polar contact between Arg87 in the mask and Glu4 in the miniprotein core (Figure 4a). In addition, Tyr82 on the mask and Trp52 in the mb core interact through pi stacking. Beyond these interactions, binding between the mask and the miniprotein is primarily hydrophobic. Although the interface between the miniprotein binder and EGFR includes 10 polar amino acid residues out of 20 residues involved in the interaction (Figure 1g), the alpha helical conformation of the mask can penetrate into the cleft between the two interacting helices. Hydrophobic residues on the mask such as Trp75, Leu83, Leu86 and Leu90 take advantage of the core of the miniprotein binder, which is rich in leucine and aromatic residues, to exclude the solvent and generate a compact four helical bundle (Figure 4a and 4b). The hydrophobic inner side of the helical mask contrasts with the highly polar solvent-exposed side, which is composed of arginine and glutamate residues (Figure 4c). The increase in overall hydrophilicity due to mask covering the hydrophobic residues at the mb interface binding with EGFR (Figure S9) could explain the higher solubility we have observed in mb_M3 with respect to EGFRn_mb, which facilitates protein concentration and handling.

**Figure 4.**
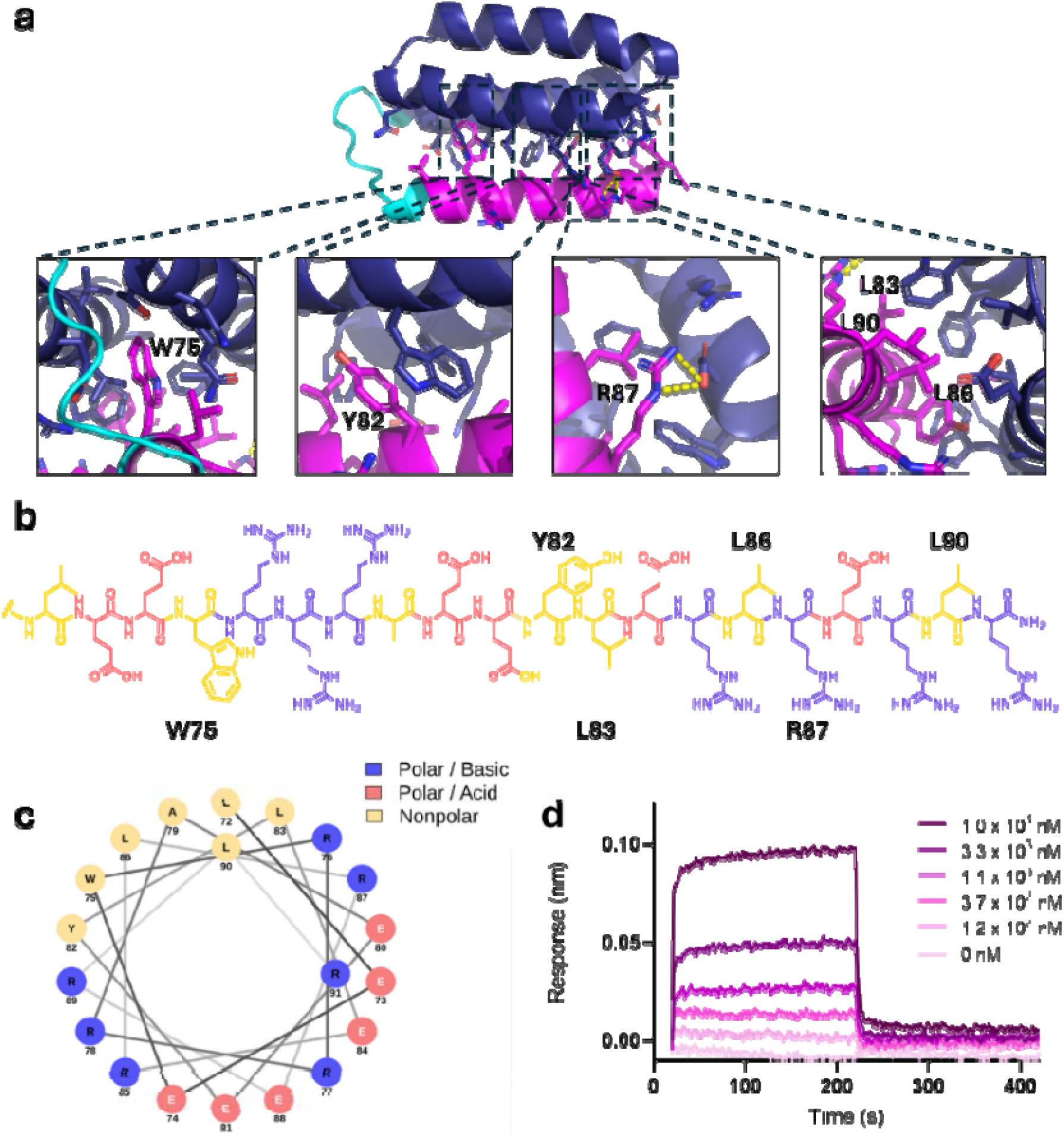
Analysis of the interaction between M3 and EGFRn_mb. a) Predicted structure of mb_M3 with AlphaFold 3. Mb core is shown in blue, linker in cyan, and mask in magenta. Residues on the mb or the mask that are within 5 Å from each other are represented as sticks in magenta or in blue, respectively. Polar interactions between the mb and the mask are shown in yellow. In panels, close view of mask residues interacting with miniprotein core. b) Amino acid sequence of the M3 mask, with nonpolar residues pale wheat color, polar acidic residues in pale red, and polar basic residues in blue. c) Helical wheel representation of the M3 mask illustrates the hydrophobic face binding the mb and the solvent-exposed hydrophilic face with polar charged residues. d) Biolayer interferometry sensograms showing binding of the M3 mask to EGFRn_mb. EGFRn_mb (mb) was immobilized on biosensors and exposed to increasing concentrations of M3. The calculated K_D_ is 5 ± 2 μM.

To assess the affinity of the M3 mask for the miniprotein binder, we produced the mask using Fmoc/tBu solid-phase peptide synthesis (Figure S10). We immobilized the miniprotein binder on a BLI biosensor and studied the binding of the mask. The calculated K_D_ of the mb for M3 is 5 ± 2 μM (Figure 4d). Although this binding affinity is 10-100 times lower than previously reported scFv-based affinity masks,^[10]^ it displays an optimal balance between outcompeting the receptor when covalently tethered to the miniprotein binder and enabling rapid diffusion of the mask upon protease cleavage.

### Light activation of the masked miniprotein binder

Given the high blocking efficiency of the tethered mask, we set out to explore whether we could make the miniprotein binder responsive to a different stimulus. We selected light as an alternative activation cue since, in contrast to proteases that rely on a local tissue expression, activation with light enables high temporal and spatial external control. However, while protease-cleavable linkers can be genetically encoded, light-cleavable linkers require building blocks that cannot be incorporated using the cell translational machinery. Therefore, we devised a chemogenetic approach to install the synthetic mask and generated a “photocaged” miniprotein binder, mb_PhM3 (Figure 5a). This strategy is based on encoding a cysteine on the miniprotein that enables the site-specific incorporation of the mask with a photo-sensitive peptide linker connecting it to the miniprotein.

**Figure 5.**
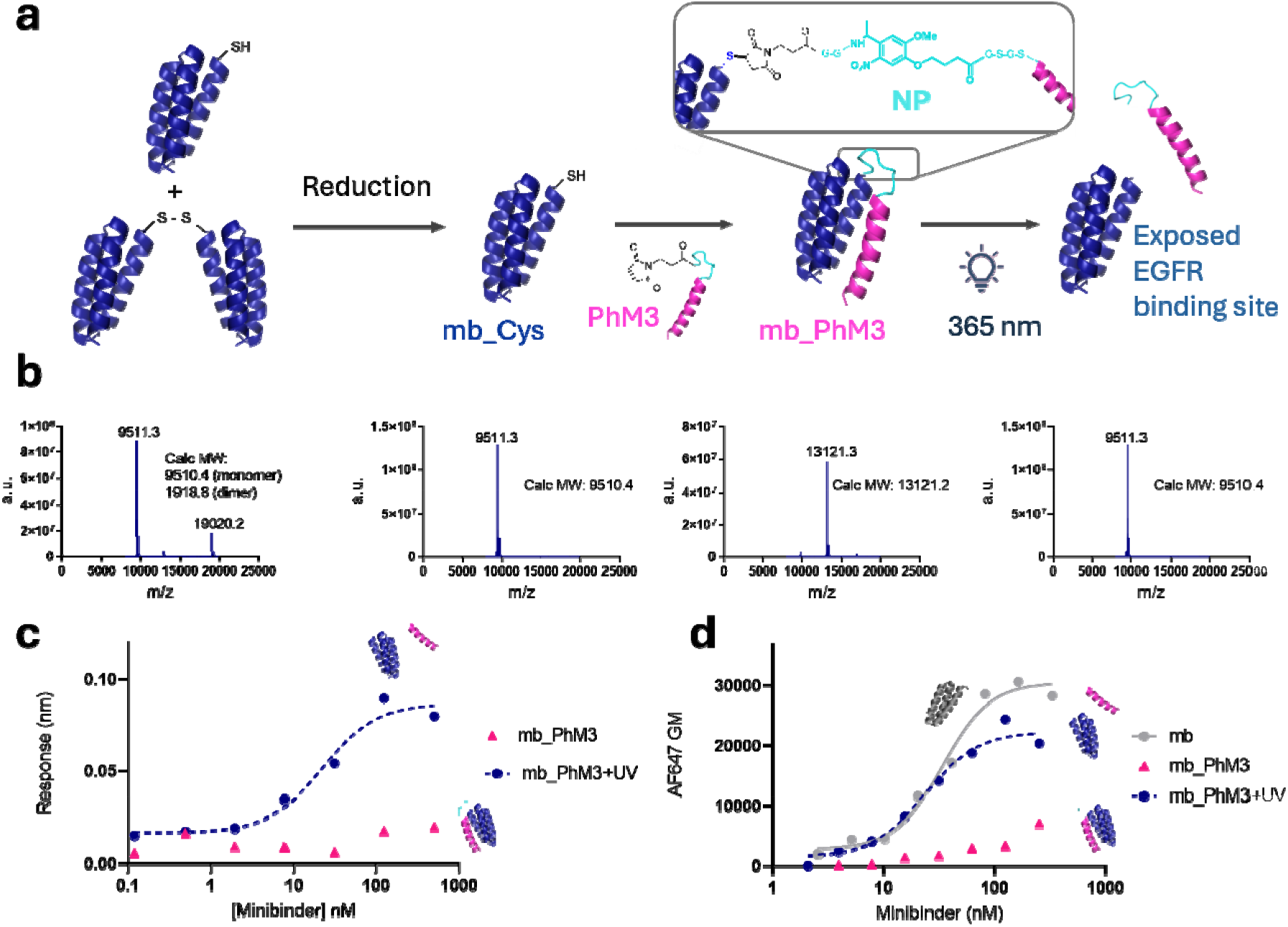
Photocaged miniprotein binder generation and binding assessment. a) Workflow scheme for production of the photo-activatable miniprotein. A partially dimerized miniprotein with a cysteine residue is reduced with TCEP to give mb_Cys in its monomeric form. The thiol group of mb_Cys is reacted with a maleimide-bearing photo-cleavable M3 mask (PhM3) to form mb_PhM3, which is cleaved upon UV light irradiation (365 nm). 3-(Maleimido-1-yl)propanoic and 4-(4-(1-aminoethyl)-2-methoxy-5-nitrophenoxy)butanoic acid are abbreviated as Mal and NP, respectively. b) Mass spectra showing, from left to right: mb_Cys as a mixture of dimeric and monomeric forms, reduced monomeric mb_Cys, photo-activated miniprotein (mb_PhM3), and mb_PhM3 after UV light activation with 90% of unmasked mb. c) Binding of mb_PhM3 to EGFR before and after photo-activation as measured by biolayer interferometry. d) Flow cytometry analysis of mb_PhM3, before and after light activation, binding to EGFR on A-431 cells.

The photo-sensitive linker bears a 4-(4-(1-aminoethyl)-2-methoxy-5-nitrophenoxy)butanoicacid moiety (NP) that can be cleaved efficiently upon irradiation with UV-light (365 nm) (Figure 5a). This moiety has been extensively applied in preclinical settings given the low toxicity of light at 365 nm.^[33]^ Glycine and Serine residues were added at each side to match the length of the protease-cleavable linker and provide sufficient flexibility for the mask to bind effectively with the miniprotein interacting surface (Figure S11). The photo-cleavable M3 mask was synthesized by Fmoc/*t*Bu solid phase peptide synthesis (Figure S12). To conjugate the mask to the miniprotein binder, we encoded a cysteine at the C-terminus of EGFRn_mb, and, to increase expression, we added a glycine residue as the terminal residue (Figure S13). Since we detected using reverse-phase HPLC coupled to mass spectrometry (LC-MS) that the cysteine-bearing miniprotein partially dimerized due to the formation of a disulfide bond (Figure 5a, 5b and S14), we added TCEP to generate the monomeric form with the free thiol (Figure 5a, 5b and S15). Subsequently, an excess of photo-cleavable M3 mask was added to ensure all miniproteins were masked and the quantitative reaction was confirmed (Figure 5a, 5b and S16). The mass spectrum indicated a systematic complete hydrolysis of the maleimide ring, which is reported to enhance the stability of the linker (Figure S17).^[32]^ The miniprotein binder was purified using nickel-NTA magnetic beads to maximize protein recovery. Purity of the photocaged miniprotein mb_PhM3 was confirmed by the presence of a single peak in a RP-HPLC UV chromatogram (Figure S18).

With the photocaged miniprotein binder in hand, we set out to prove that binding could be rescued upon light irradiation (Figure 5a). mb_PhM3 was irradiated with UV light at 365 nm for 2 h. LC-MS shows over 90% release of the mask (Figures 5b and S19). The mass spectrum displays a main peak with a mass of 9811 Da, which matches the expected mass of the cleaved mb_PhM3, while the peak of the masked miniprotein at 13121 Da is barely observable. We next proved that the miniprotein binder with the photo-cleavable mask has negligible binding to the EGFR receptor immobilized on the BLI biosensor (Figure 5c). The affinity decrease generated by this mask is over 2 orders of magnitude, only slightly lower than the protease-sensitive version. Upon activation with light, binding is fully recovered.

The same experiment was repeated by incubating the miniprotein binder prior or after irradiation with cells expressing high levels of EGFR at 4°C. Flow cytometry analysis of ight the cells showed that the EC_50_ of the parental miniprotein binder is fully recovered (Figure 5d). The fact that both photo-cleavable and protease-cleavable masks provide a highly efficient masking despite their different physicochemical properties indicates that the linker has little effect on the masking capacity of the binding sequence.

## CONCLUSION

In this study we have developed a method to reversibly inactivate miniprotein binders with a minimal mask designed *de novo* and have proved its applicability on a miniprotein that antagonizes EGFR. We have shown that a combination of protein design tools enables the extension of the C-terminus of a miniprotein to cover the binding interface with a masking moiety. This computational design workflow proved to be highly efficient since the six designs produced were able to substantially decrease the binding affinity of the miniprotein for EGFR. Affinity reduction was of two orders of magnitude in 5 designs and over 1000 times for the best design, M3. Furthermore, binding affinity was quantitatively recovered for all mask designs tested after protease cleavage of the mask, proving that inactivation is fully reversible. Thus, the single digit micromolar affinity of the M3 mask for the miniprotein binder enables efficient blockage when tethered but can be effectively outcompeted by EGFR upon cleavage with proteases. We have also shown that reversible inactivation of this miniprotein enables control over downstream EGFR signaling in cells. Finally, site-specific conjugation of a synthetic mask allows to easily render the reversibly inactivated miniprotein responsive to other stimuli such as light. Overall, we have engineered the first *de novo* designed masked miniprotein binders with masks responsive to stimuli such as proteases or light. Furthermore, we have shown for the first time that peptide masks that reversibly inactivate proteins can be computationally designed from scratch with a highly efficient workflow. Our strategy opens the doors to accelerating the development of affinity-based masks for other proteins with therapeutic potential rendering them conditionally active to any stimulus.

## NOTES

The authors declare no conflict of interest.

Supplementary information will be made available upon publication in a journal.

## ACKNOWLEDGEMENTS

EMBO and Fulbright Ruth-Lee Kennedy travel grants to B.O-S. enabled starting the project. We acknowledge support from the European Research Council (ERC) under the European Union’s Horizon Europe research and innovation program (grant agreement No. 101077370) and from MCIN/AEI/10.13039/501100011033 (PID2023-151988OB-I00), the European Social Fund Plus (ESF+). The project that gave rise to these results received the support of fellowships from the “la Caixa” Foundation (ID 100010434). The fellowship codes from the INPhINIT and Junior Leaders programs to M.E-R. and B.O-S. are LCF/BQ/DR23/12000019 and LCF/BQ/PR21/11840002, respectively. C.M., R.L., and C.D.-P., held FPI (PRE2023-002030 MCIN/AEI), FPU (FPU19/03216 MCIN/AEI), and MSCA-PF 101063066 fellowships, respectively.

## Notes

### Competing Interest Statement

The authors have declared no competing interest.

